# Using nature-based citizen science initiatives to enhance nature connection and mental health

**DOI:** 10.1101/2024.07.14.603425

**Authors:** Rachel R. Y. Oh, Andres F. Suarez-Castro, Richard A. Fuller, Michael Tervo, Kevin Rozario, Birte Peters, Shawan Chowdhury, Julia von Gönner, Martin Friedrichs-Manthey, Ambros Berger, Tracy Schultz, Angela J. Dean, Ayesha Tulloch, Aletta Bonn

**Author notes:** **Correspondence:** Rachel R. Y. Oh.

## Abstract

The global rise in mental health issues underscores the critical importance of assessing the mental health benefits of engaging with nature. Beyond their primary aim of involving citizens in scientific data collection, nature-based citizen science initiatives offer significant potential for enhancing conservation (e.g. connection to nature) and human health and wellbeing (e.g. emotions, depression, stress, anxiety) outcomes. However, the effectiveness of various types of initiatives in achieving specific health outcomes remain unclear. This study evaluates changes in eight nature connection and health and wellbeing outcomes before and after participation in five such initiatives, specifically three dimensions of connection to nature (Self, Experience and Perspective) measured by the Nature Relatedness scale), and mental health outcomes—depression, stress and anxiety—measured by the DASS-21 scale; along with positive and negative emotions assessed using the Scale of Positive and Negative Experience (SPANE). We found that participants generally reported improvements across all outcomes, but only positive emotions and reduced anxiety and stress symptoms were significantly enhanced. This suggests that short-term nature-based interventions are effective in boosting emotions and alleviating anxiety and stress, though significant changes in nature connection and depression may require prolonged or more intensive engagement. We also found that the Queensland Trust for Nature initiative, characterized by its extended duration and social interactions, emerged as a particularly effective model in achieving synergies between biodiversity and human health goals. Consequently, we advocate for the reimagining of these nature-based initiatives as integral components of broader health-promoting strategies. By aligning citizen science efforts with health promotion frameworks, we can amplify the utility and impact of nature-based citizen science initiatives, enhancing their contributions to scientific knowledge, nature conservation and human health.

## 1 Introduction

Citizen science initiatives aimed at crowd-sourcing biodiversity data are rapidly expanding in both scale and scope. They involve public participation in scientific research and knowledge production, and have evolved into a well-developed and valued approach with global applicability across various scientific disciplines (Fraisl et al., 2020; Kullenberg & Kasperowski, 2016). Rising literacy levels and the widespread adoption of information technology infrastructure have empowered individuals to contribute meaningfully to scientific knowledge creation (Aristeidou & Herodotou, 2020). As such, citizen science data is now a leading source of biodiversity information (Fritz et al., 2019) as they help bridge important temporal and spatial knowledge gaps (Bradter et al., 2018; La Sorte & Somveille, 2020). In many regions of the world, they are used to monitor biodiversity changes (Forister et al., 2021), inform on-the-ground species management strategies (Beninde et al., 2023), and to understand ecological processes and species interactions (Groom et al., 2021). For example, bird biodiversity data from the citizen science initiative eBird has been used to model species’ distributions, abundances, and how they change over time (Fink et al., 2020).

Beyond contributing valuable biodiversity data, engagement in biodiversity citizen science initiatives can additionally enhance participants’ personal wellbeing and improve conservation outcomes by amplifying environmentally protective behaviors. Surveys of citizen science participants suggest that involvement in these initiatives enhances their personal wellbeing through enjoyment of the activity, improved scientific literacy and connecting with like-minded individuals (Kelemen-Finan et al., 2018; Peter et al., 2021). Participants often exhibit heightened awareness of environmental issues (Peter et al., 2021) and develop a stronger connection to nature (Nisbet et al., 2009), which is an important driver of active participation in conservation efforts (Whitburn et al., 2020). Individuals who are strongly connected to nature also derive significant health and wellbeing benefits from exposure to nature (Lackey et al., 2021; Oh et al., 2021).

From a health perspective, biodiversity citizen science initiatives represent a form of nature-based health intervention. These are programs, activities or strategies designed to enhance health and wellbeing (Shanahan et al., 2019) and are centered around natural elements such as vegetation and water bodies. They generally encompass a diverse range of activities from the development of community gardens, to sea swimming initiatives and wilderness programs (Hunter et al., 2019). In fact, the growing interest in nature-based interventions is underscored by the global mental health crisis (Van Den Bosch & Ode Sang, 2017), and the substantial annual healthcare cost savings provided by protected nature areas worldwide estimated at US$6 trillion (Buckley et al., 2019). Urban and public health administrations are beginning to acknowledge the importance of proximity to, and interaction with, natural environments as proactive health interventions for populations (Maller et al., 2006).

Given that most biodiversity citizen science initiatives require outdoor participation for data collection, they inherently foster health-promoting behaviors in nature such as physical exercise through walking and hiking (Biddle et al., 2019; Warburton et al., 2006) or help reduce psychological stress and enhance cognitive function (Jimenez et al., 2021). The inherently social and collaborative nature of these initiatives further fosters social connections, thereby reducing social isolation―these are key factors for strengthening long-term mental resilience and health (Jordan et al., 2011). As such, biodiversity citizen science initiatives present a critical opportunity for healthcare systems to offer a broader suite of lifestyle intervention (Britton et al., 2020). Consequently, there is a rapidly growing interest among conservation organizations (Carr & Hughes, 2023) and global biodiversity and human health policy platforms (IPBES, 2019) in leveraging nature-based citizen science initiatives for broader impacts as incorporating health and well-being as a recruitment strategy in citizen science initiatives can serve to attract a broader subset of society, thereby enriching the dataset available for conservation efforts.

There is a pressing need to comprehensively investigate the emergence of such benefits across various nature-based citizen science initiatives as they have a weak evidence base (Pocock et al., 2023). We do not know if engagement in nature-based citizen science initiatives are linked to enhanced health outcomes and behavioral change, and if yes, which types of engagement are most effective in achieving which targeted conservation or health outcomes. For instance, would participating in a citizen science initiative for one hour per week be more effective in reducing stress compared to engaging for 30 minutes twice a week? In this study, we employed a before-and-after design to investigate which type of nature-based citizen science initiatives are associated with enhanced nature connection and mental health outcomes. We do so across five nature-based citizen science initiatives that varied in the duration, frequency and intensity of participants’ engagement and the ecosystems or taxa studied. We assessed whether any positive changes could be attributed to project-specific and individual-specific factors. Additionally, we highlight the biodiversity data contributions from these initiatives to demonstrate their primary role in data acquisition towards informing conservation efforts. Understanding how engagement in citizen science initiative can enhance nature connection and health outcomes is crucial for the effective design of initiatives that achieve these benefits alongside their primary objectives of collecting biodiversity data.

## 2 Materials and Methods

### 2.1 The five nature-based citizen science initiatives

We compared five nature-based citizen science initiatives (Figure 1), encompassing four initiatives based in Germany (FLOW―Exploring Streams, Creating Knowledge Together: www.flow-projekt.de; Freshwater Detectives; VielFalterGarten―Many Butterfly Gardens: www.vielfaltergarten.de; and Pflanze KlimaKultur!―Plant Climate Culture!: www.pflanzeklimakultur.de) and one from Australia (QTFN―Queensland Trust for Nature: https://qtfn.org.au/). Within each initiative, each data collection event varied in frequency and average duration, ranging from 15 minutes (1 – 3 days per week) to 48 hours (once annually). Data collection for all initiatives were conducted individually, except for FLOW and QTFN which were group-based. While these initiatives aim to collect biodiversity and environmental data using standardized methods across diverse ecosystems such as unmanaged nature, freshwater streams, canals and urban greenspaces, each initiative is unique in how the collected data is used to inform conservation efforts (Figure 1). Data collected from the Germany-based initiatives were reported directly to the researchers via tailored apps and recording schemes, while the citizen science records from the Australian-based initiative were reported directly to <u>iNaturalist</u>.

**Figure 1:**
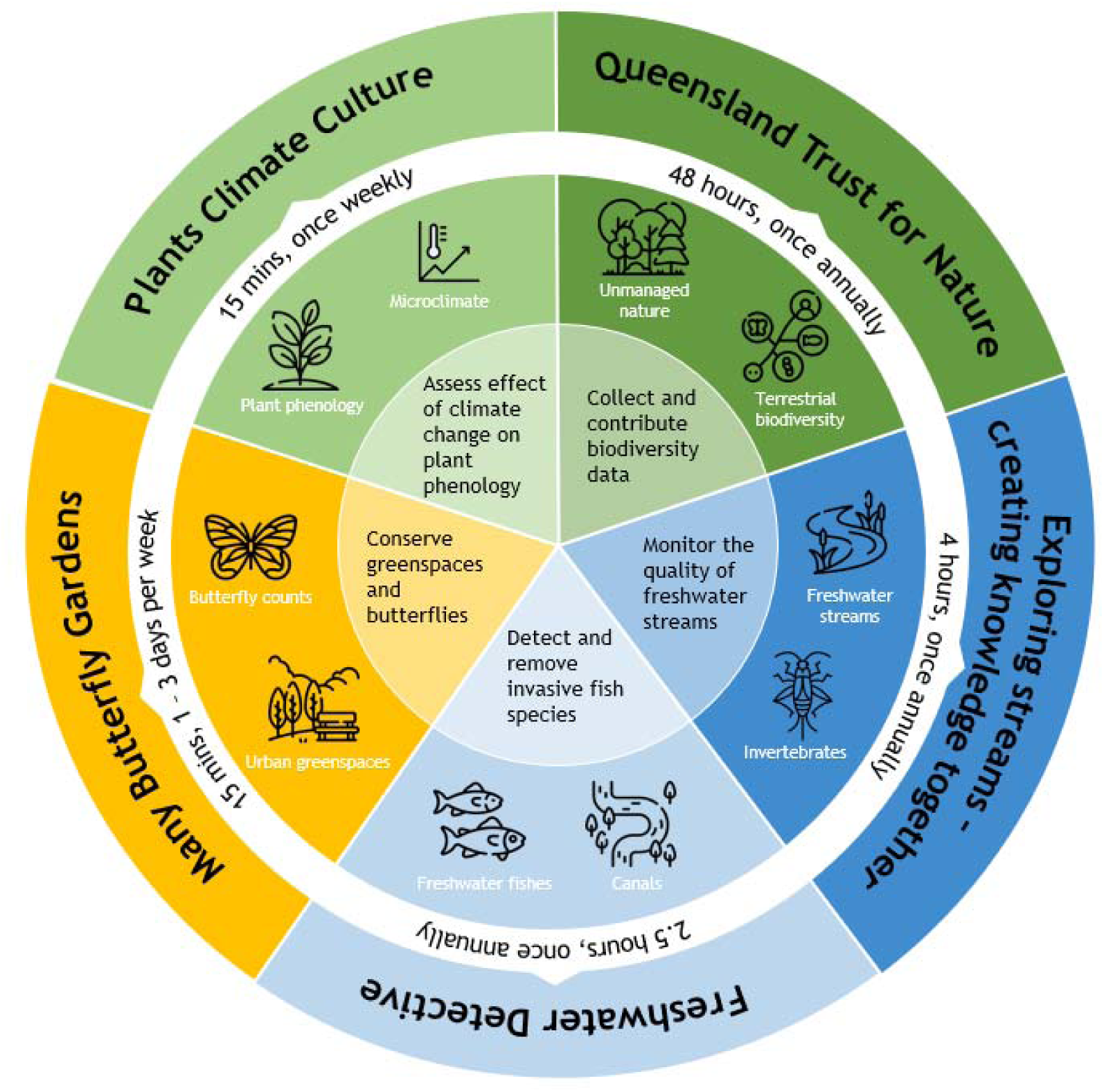
An overview of the five nature-based citizen science initiatives and each, encompassing four initiatives based in Germany (FLOW―Exploring Streams, Creating Knowledge Together; Freshwater Detectives; VielFalterGarten―Many Butterfly Gardens; and Pflanze KlimaKultur!―Plant Climate Culture!) and one from Australia (QTFN―Queensland Trust for Nature). Each data collection event (depicted by the white ring) varied in frequency and average duration, ranging from 15 minutes (1 – 3 days per week) to 48 hours (once annually). Each initiative is unique in the ecosystem and taxa studied, which varied from freshwater streams and canals, to unmanaged and urban greenspaces. Each initiative also varied in how the collected data is used to inform conservation efforts (innermost circle).

### 2.2 Study Design and Participant Recruitment

We administrated surveys across five distinct nature-based citizen science initiatives in Australia and Germany from March to November 2023 to adults aged 18 years and above. Surveys were delivered to participants prior to participation, and immediately after their participation. Each citizen science initiative varied in terms of how often (“frequency”) and how long (“duration”) participants collected biodiversity data, as well as the ecosystems wherein data was collected (see Figure 1 for specific details on each citizen science initiative). In total, we received 278 surveys from the following initiatives – QTFN (63 surveys); FLOW (72 surveys); Freshwater Detectives (42 surveys); VielFalterGarten―Many Butterfly Gardens (8 surveys) and Pflanze KlimaKultur!―Plant Climate Culture! (93 surveys). Participants were recruited through public promotion by research and initiative partners, and also involved circulation via social media. There was no pre-requisite for participants to participate in these initiatives. Prior to survey commencement, all participants provided informed consent by ticking a consent box. This research was conducted in accordance with the Declaration of Helsinki, ethical guidelines provided by the University of Griffith Institutional Human Research Ethics (Reference Number: 2023/190) and the UFZ Datenschutz (Data Protection Committee) at the Helmholtz-Zentrum für Umweltforschung GmbH – UFZ (Approval: 08132024).

### 2.3 Response Variables (Nature Connection and Mental Health and Wellbeing)

We administered the same survey across all five citizen science initiatives to assess a total of 11 self-reported outcomes pertaining to individuals’ connection to nature, and their mental health and wellbeing. The full surveys administrated before and after participation can be found in the Supplementary Materia (see Appendices A and B), and we elaborate on each of these outcomes below.

#### 2.3.1 Connection to nature

Participants assessed their connection to nature using the nature-relatedness scale (Nisbet et al., 2009). Participants were invited to rate a set of 21 statements using a five-point Likert scale ranging from 1 (disagree strongly) to 5 (agree strongly). This scale captures three dimensions of an individual’s relationship with nature―NR-Experience (experiential), which indicates physical familiarity with, and attraction to, nature (e.g., “I enjoy being outdoors, even in unpleasant weather”; NR-Self (affective), which assesses how strongly one identifies with nature (e.g., “My relationship to nature is an important part of who I am”); and NR-Perspective (cognitive), which indicates one’s personal attitude and behaviors towards the environment (e.g., “Conservation is unnecessary because nature is strong enough to recover from any human impact” 2009).

While a composite score across all 21 statements provides an overall measure of nature connection, we analyzed these dimensions separately to understand how participation shaped three different elements of connection, such as physical proximity to nature (e.g. by being outside and physically immerse in nature) or a deeper psychological connection (e.g. a sense of feeling a part of nature) (Butler et al., 2024). Moreover, a meta-analysis has suggested that while physical connection with nature enhances short-term psychological connection, it is the emotional connection that has a stronger influence on conservation behaviors (Barragan-Jason et al., 2023).

#### 2.3.2 Mental health

We assessed the severity of depression, anxiety and stress symptoms experienced before and during engagement using the Depression, Anxiety and Stress Scale (DASS-21), a standardized global self-reporting questionnaire (Lovibond & Lovibond, 1995). Participants rated 21 statements on a four-point Likert scale (‘0 – Not applicable to me, ‘1 = Applicable some of the time’, ‘2 = Applicable for a good part of time’, and ‘3 = Applicable for most of the time’). Within these statements, three sets of 7 statements measured symptoms of depression such as feelings of disinterest and inertia, anxiety such as situational anxiety and subjective experiences of anxious effects and stress such as having difficulty relaxing, nervous arousal, irritability.

In addition to the DASS-21, participants reported their levels of anxiety, stress and ability to concentrate via three visual analog scales. These scales are depicted as horizontal lines anchored by “not at all” on one end, and “very much” on the other, and were chosen for their simplicity and ability to capture immediate perceptions of mental states. While validated scales are the gold standard in psychology for their reliability (i.e. produce consistent results across different contexts) and validity (i.e. accurately measure the constructs they are developed to measure), they typically feature a larger number of statements to capture nuanced variations in outcomes and therefore could be less responsive at detecting subtle short-term changes (García-Pérez & Alcalá-Quintana, 2023). We therefore trialed visual analog scales to measure mental wellbeing (Elo et al., 2003) for two reasons (i) we expected rapid fluctuations in outcomes before and after participation; and (ii) we aimed to have a concise survey to minimize participant fatigue.

#### 2.3.3 Wellbeing (Positive and negative affect)

We assessed participants’ broad range of experiences and feelings using the Scale of Positive and Negative Experience (SPANE; (Diener et al., 2009)). Participants were invited to indicate the frequency with which they experienced six positive (e.g. happy, contented) and six negative emotions (e.g. angry, sad) before and immediately after participating in the citizen science initiative. There were five response options for each emotion, ranging from “very rarely or never” to “very often or always. We chose the SPANE over the Positive and Negative Affect Scale (PANAS; (Watson et al., 1988)) as it offered a more balanced range of emotions in terms of arousal, is sensitive to differences across cultures and contexts (Diener et al., 2009).

### 2.4 Predictor Variables (Socio-demographic variables)

We collected socio-demographic covariates including age, gender (male, female, non-binary), number of hours worked per week and number of hours spent in greenspaces in the past week, as these have been tied to mental health outcomes and experiences of nature in previous studies (Elliott et al., 2023; Shanahan et al., 2016; White et al., 2023).

### 2.5 Statistical Analyses

We conducted all analyses using R version 4.4.1 (R Core Team, 2024). After excluding incomplete surveys, our analyses included data from 253 completed surveys, representing 91% of 278 received surveys. This comprised 49 surveys from QTFN, 68 surveys from FLOW, 35 surveys from Freshwater Detectives, 8 surveys from VielFalterGarten―Many Butterfly Gardens and 93 surveys from Pflanze KlimaKultur!―Plant Climate Culture!.

We constructed a total of 11 global models to examine the correlation between each outcome and potential predictor variables. We used cumulative link mixed models (CLMM; ordinal package; Christensen, 2023) for eight outcomes: three dimensions of connection to nature (NR Self, NR Experience, NR Perspective); depression, stress and anxiety as measured using the DASS-21 scale; and positive and negative affect measured using the SPANE. We do so as the response variable was ordered, as it was specified as the change in rating for each statement, and was computed as the rating of statement after participation minus rating of statement before participation. This change could range from −4 to 4 for the dimensions of nature relatedness (NR Self, NR Experience, NR Perspective); −3 to 3 for symptoms of depression, anxiety and stress measured using the DASS-21, and −4 to 4 for positive and negative affect. An improvement after engagement in the nature-based citizen science initiatives was defined as an enhanced connection to nature, a reduction in symptoms of depression, anxiety and stress, fewer negative emotions, more positive emotions.

We used generalized linear models for the three visual analog scales (anxiety, stress and ability to concentrate) as the response variables were continuous. Similarly, this change could range from −10 to 10, and an improvement after engagement in the nature-based citizen science initiatives was defined as a reduction in symptoms of anxiety and stress, and stronger concentration.

For all 11 models, the predictor variables specified were project, age, gender, number of hours worked and number of hours spent in greenspaces, which are summarized in Table 1. All continuous predictors were standardized. For the CLMMs, each respondent was additionally specified as a random intercept. We assessed the goodness-of-fit for CLMMs and GLMs using the Hessian condition number (Christensen, 2023) and residuals of fitted models respectively. Prior to all analyses, we assessed multicollinearity in each global model using the vif function from the car package (Fox & Weisberg, 2023), and found no such issues (VIF < 3).

**Table 1:** Description of the predictor variables specified in each of the 11 global models where response variables were: three dimensions of connection to nature (NR-Self, NR-Experience, NR-Perspective); depression, stress and anxiety as measured using the DASS-21 scale; stress, anxiety and ability to concentrate as measured using the visual analog scale, and positive and negative affect.

| Variable | Description |
| --- | --- |
| Project (categorical) | Five different projects – QTFN—Queensland Trust for Nature; FLOW—Exploring Streams, Creating Knowledge Together; Freshwater Detectives; VielFalterGarten—Many Butterfly Gardens; and Pflanze KlimaKultur!—Plant Climate Culture! |
| Age (linear) | Participants provided their age in years. |
| Gender (categorical) | Female, male, non-binary. |
| Number of hours worked per week (linear) | The number of hours the participant worked in an average week. |
| Duration of greenspace visits (linear) | Self-reported average number of hour(s) spent during each visit to public outdoor greenspaces in the week prior to participating in the initiative. |

As it was possible that a different statistical treatment of the response variables (from ordered to continuous) might result in a different set of significant predictors, we then conducted an additional analysis that used generalized linear models instead of CLMMs. We aggregated the responses for each outcome to form a continuous score that provided a measure of an individual’s connection to nature, mental health and wellbeing (as per Diener et al., 2009; Lovibond & Lovibond, 1995; Nisbet et al., 2009). We report these results in the Supplementary Material (Figure S2) given that the significant predictors and direction of relationships with response variables of generalized linear models were generally consistent with the CLMMs.

### 2.6 Biodiversity Measures

For each citizen science initiative, we also collected data on species richness and abundance documented by the participants. This encompassed species from terrestrial and freshwater ecosystems in Australia and Germany, and plant phenology from Germany (Figure 1).

## 3 Results

While we constructed a total of 11 global models, the results presented here are derived from the eight CLMMs that demonstrated good model fit. They represent the following outcomes: three dimensions of connection to nature, three aspects of mental wellbeing (depression, stress and anxiety symptoms) measured using the DASS-21 scale, and positive and negative affect. The results from the three models based on data collected via the visual analog scales are not reported due to poor model fit.

### 3.1 Overall change in nature connection and mental health and wellbeing

Across all eight outcomes related to nature connection and mental health and wellbeing, the highest percentage of change (in ratings) was observed for negative and positive affect, as well as symptoms of stress (Figure 2, but see Figure S1 for a detailed distribution of change in ratings for each citizen science initiative). 55.8% of participants reported fewer negative affect, while 42% reported greater positive affect (Figure 2). 47.7% of participants reported lesser symptoms of stress (Figure 2). These results indicate that participants experienced improved mental health and wellbeing outcomes after participating in the nature-based citizen science initiatives. Regardless of whether the majority of participants reported a change in mental health and wellbeing, there were consistently more improvements than worsened outcomes, ranging from 1.3 times more improved versus worsened outcomes for NR-Self and NR-Experience (which also had the highest percentage of outcomes categorized as worsened: 13.1% and 12.3%, respectively) to 11.6 times more reports of improved versus worsened outcomes for negative affect. Anxiety exhibited the lowest percentage of outcomes classified as worsened (3.2%).

**Figure 2:**
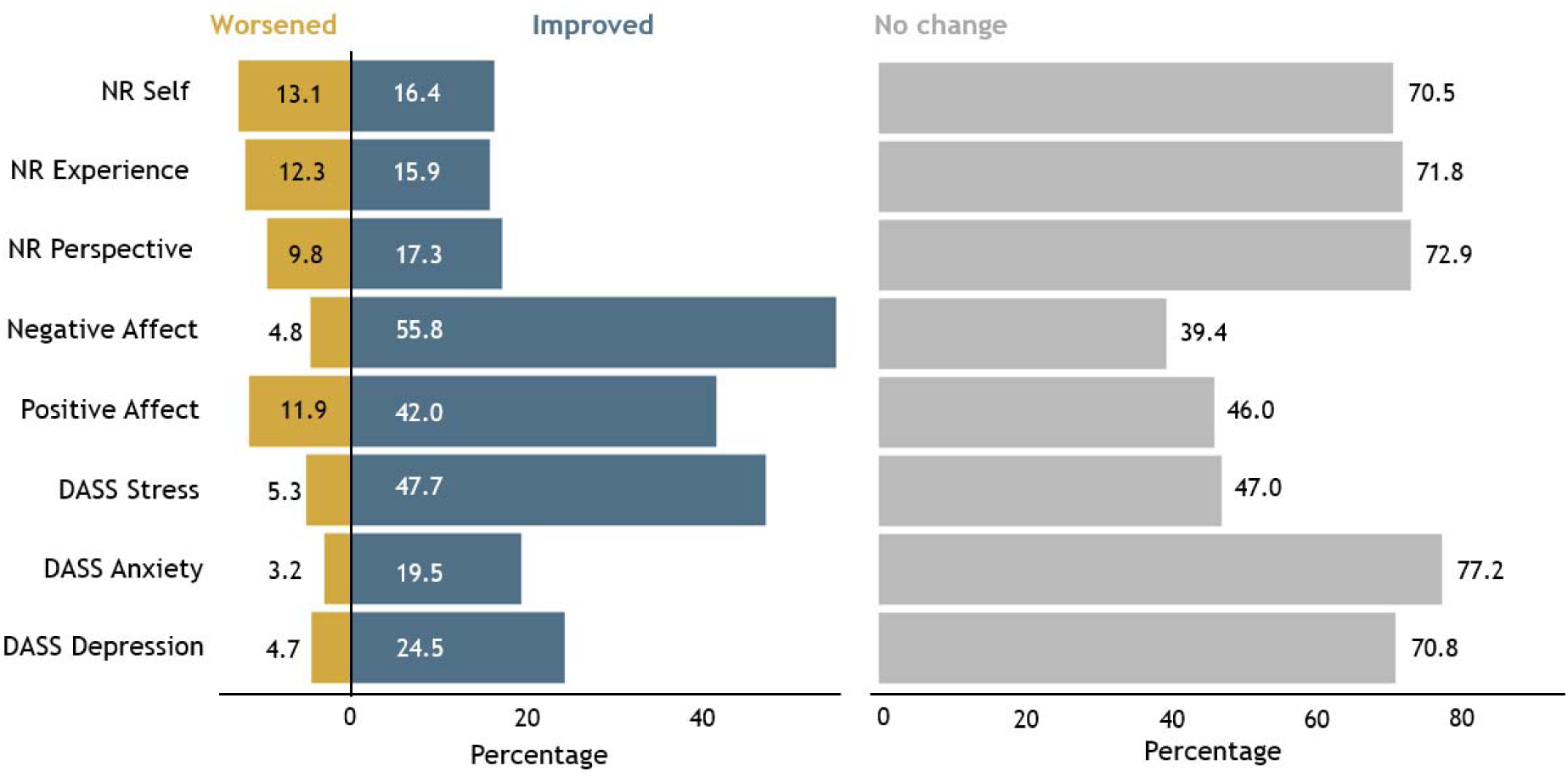
The percentage change in ratings for each outcome was categorized as improved (colored blue), worsened (colored gold), or remained unchanged (colored grey). The ratings in the improved and worsened categories have been aggregated; to illustrate, changes in ratings under the DASS21 scale have been aggregated as follows: negative changes (−3, −2, −1) are categorized as “improved,” positive changes (+3, +2, +1) are categorized as “worsened,” and no change (0) is categorized as “unchanged.“

### 3.2 Relationship between citizen science participation and outcome measures

We used CLMMs to assess the relationship between participation in nature-based citizen science initiatives and nature connection and mental health and wellbeing outcomes. Overall, after controlling for socio-demographic predictor variables, we found that engagement in citizen science initiatives was not consistently associated with an enhanced connection to nature or improved health and wellbeing. Among the 11 outcomes studied, only eight models exhibited good model fit (Hessian condition number < 1 × 10^4^; Christensen, 2023). The three models incorporating data from the visual analog scales did not meet this criterion.

Of the eight well-fitted models, only four outcomes had significant predictors (Figure 3). First, a stronger connection to nature (NR-Experience) was associated with increased time spent in greenspaces (Figure 3a). Second, older participants reported greater reduction in symptoms of anxiety and stress compared to younger participants (Figure 3b). Third, participating in the QTFN initiative was positively associated with positive affect, but male participants reported lower levels of positive affect compared to female participants (Figure 3c). No predictor variables emerged as statistically significant in CLMMs where NR-Self, NR-Perspective, DASS Depression and Negative Affect were the specified response variables.

**Figure 3:**
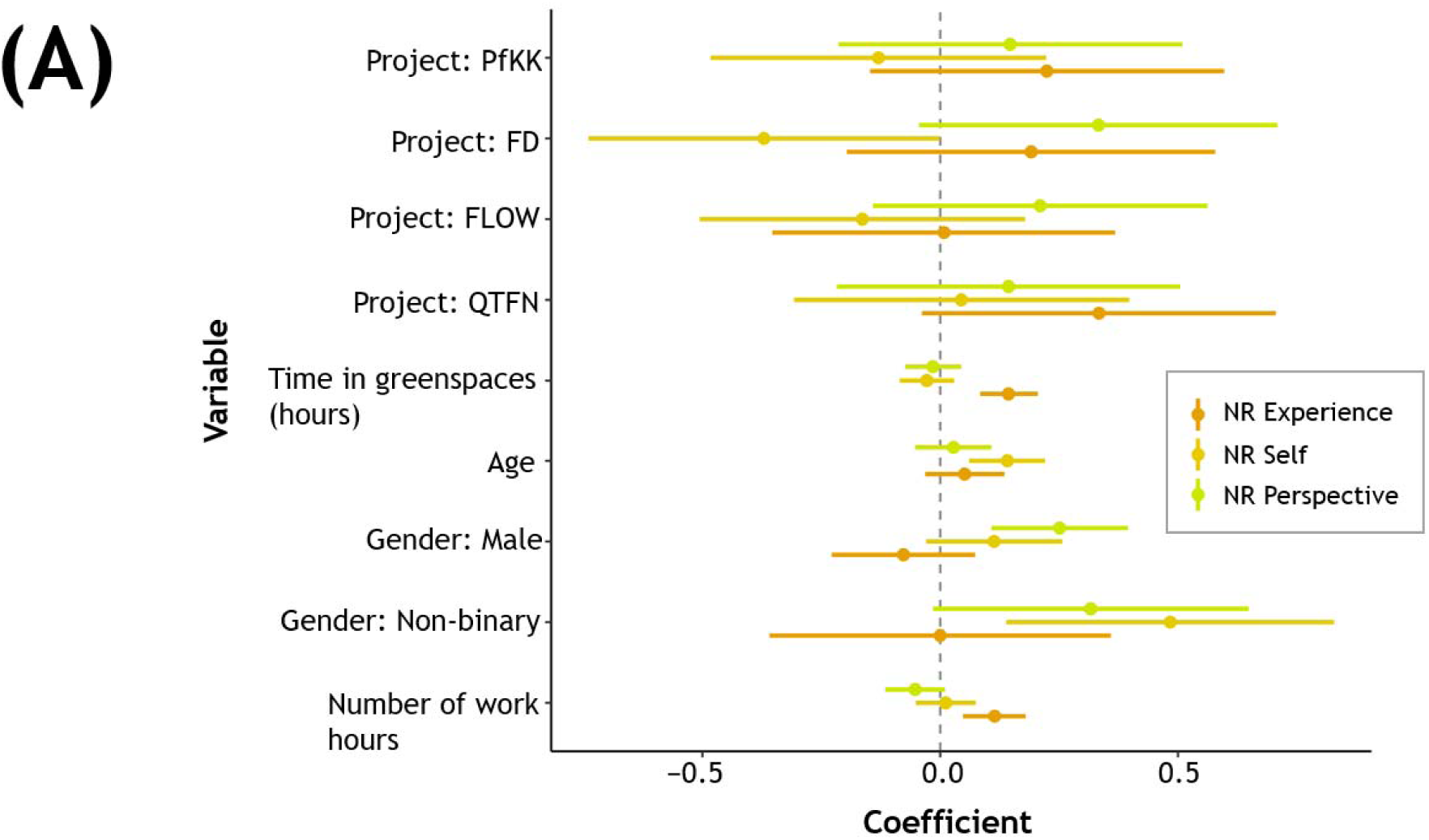
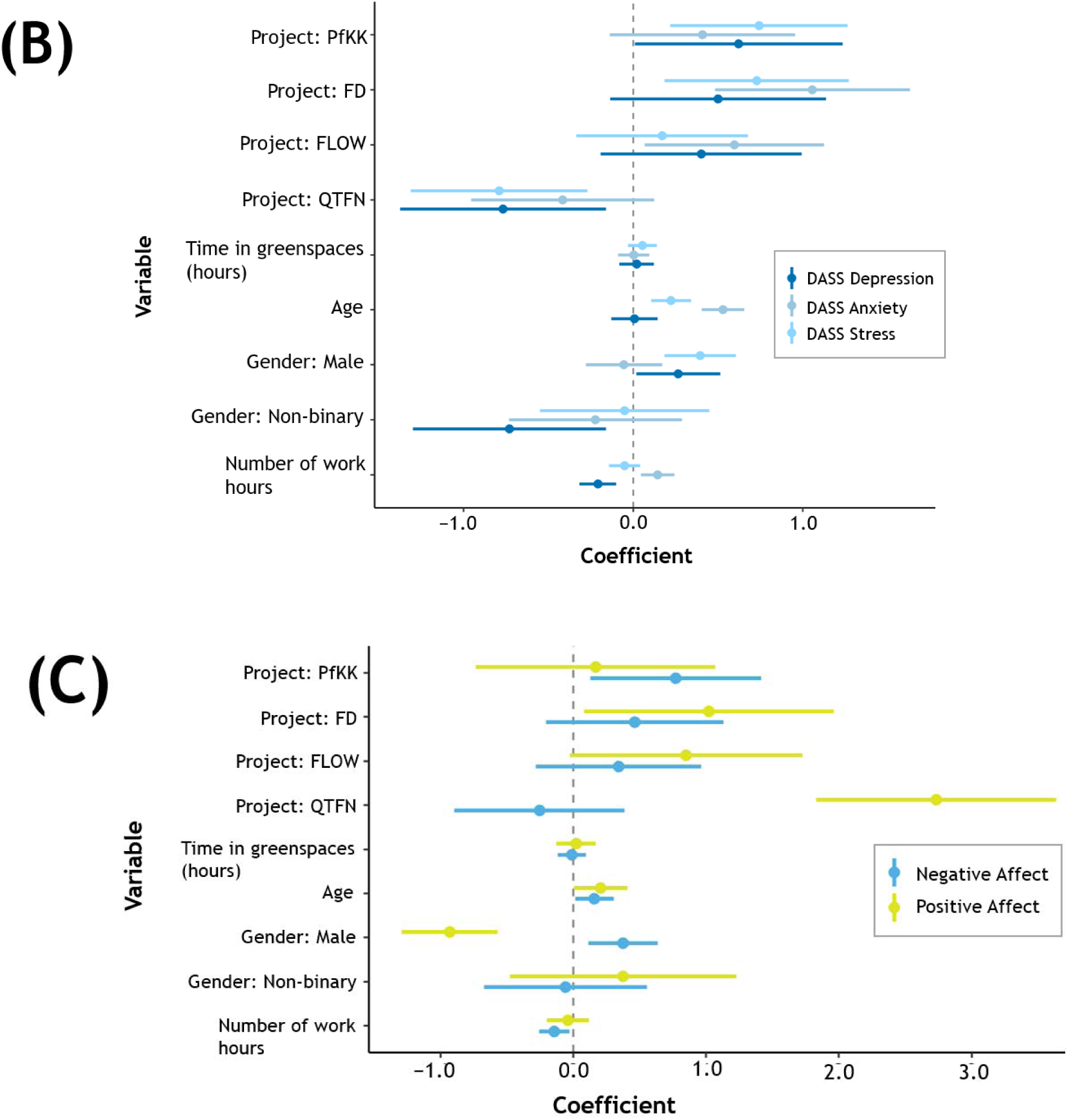
The model outputs from the CLMMs, showing the effect size and standard errors of each predictor variable for each outcome of (a) three dimensions of nature connection (NR-Experience, NR-Self and NR-Perspective; (b) depression, anxiety and stress as measured using the DASS-21; and (c) negative and positive affect. While we presented the effect size and standard errors for all predictor variables, please refer to Section 3.2 for the exact list of significant estimated coefficients (p-value ≤ 0.05). The effect size and standard errors for categorical variables are presented relative to a comparative base factor level: Project (Many Butterfly Gardens) and Gender (Female).

### 3.3 Biodiversity

As active engagement in acquiring biodiversity data was the primary goal of these citizen science initiatives, we highlight also the biodiversity data contributions from participations. When comparing the biodiversity data efforts across citizen science initiatives, we noticed substantial differences in identifying observations up to individual taxonomic levels (Figure 4a). For example, for the QTFN initiative, 72% of the observations were reported as species, whereas the FLOW initiative could only record benthic invertebrates (macrozoobenthos) at the family level and < 20% of records were at species level. For Freshwater Detectives and Pflanze KlimaKultur!―Plant Climate Culture!, only species-level information was reported (Figure 4), as the former had low levels of freshwater fish biodiversity in canals and the latter had only 11 targeted species. For the Pflanze KlimaKultur!―Plant Climate Culture! Initiative however, there existed additional ecology data such as plant phenology (e.g. months when budding, fruit ripening, leaf unfolding, aging was observed; Figure 4b) and how that could change in response to changes in urban microclimate (Figure 4b).

**Figure 4:**
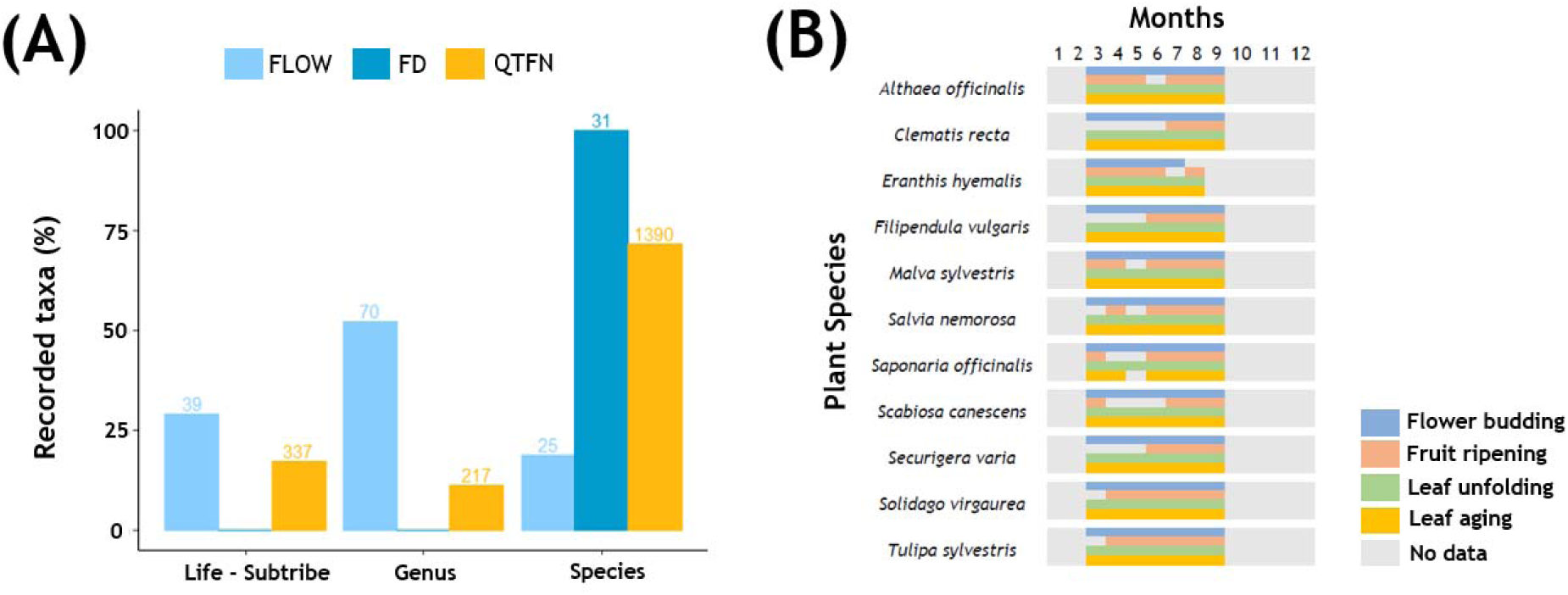
The scientific data contributed varied between the citizen science initiatives. (a) The number of observations identified across taxonomic levels of Life, Genus and Species from three citizen science initiatives (FLOW, FD and QTFN). ‘Life – Subtribe’ includes Life, Kingdom, Phylum, Class, Subclass, Order, Suborder, Infraorder, Superfamily, Clade, Family, Subfamily, Tribe and Subtribe. The number at the top of each bar represents the total number of taxa identified at that taxonomic level. (b) The four key stages of plant phenology (flower budding, fruit ripening, leaf unfolding and leaf aging) documented by participants for 11 plant species across seven months from the Pflanze KlimaKultur!―Plant Climate Culture! Initiative.

## 4 Discussion

Assessing the mental health benefits of engaging with nature is important, given the rising prevalence of mental health issues globally (Van Den Bosch & Ode Sang, 2017), and the considerable potential for healthcare cost savings through nature-based interventions (Buckley et al., 2019). This study aimed to evaluate the contributions of nature-based citizen science initiatives in enhancing human health outcomes, alongside their primary objective of engaging citizens in scientific data collection.

### 4.1 Mental health and wellbeing outcomes vary in size of response to short-term nature-based interventions

Using a consistent assessment method across five nature-based citizen science initiatives, we found that the magnitude of improvement varied across the eight measured outcomes. Participants generally reported enhanced nature connection and improved mental health and wellbeing, with the most substantial changes observed in negative and positive emotions, as well as stress symptoms (Figure 2). This indicated that changes in emotions and stress symptoms are more responsive to short-term nature-based intervention. This finding aligns with studies suggesting that emotions are preconditions to human behaviors, allowing individuals to quickly process and adapt or respond to external situations, such as fear triggering a fight, flight or freeze response (Brosch, 2021). As such, these outcomes are more responsive to short-term changes compared to other health outcomes such as depression and anxiety, which develop over time from cumulative, recurring life experiences (Kessler & Bromet, 2013). For example, symptoms of depression and anxiety persist for at least two weeks and six months respectively, durations that exceed any citizen science initiative (the longest duration of which lasted only 48 hours). Such outcomes therefore require a series of regular interventions or lifestyle changes before any significant improvement can be observed (Sarris et al., 2020; Shorey et al., 2022).

Therefore, at the individual-level, the efficacy of nature-based (health) interventions is likely maximized when (i) targeting outcomes responsive to short-term stimuli such as emotions and stress; and (ii) delivering these interventions regularly, rather than as one-off events to ensure sustained benefits over time. At the society or population level, the efficacy of such nature-based interventions is likely better suited to foster and amplify health-promoting behaviors (rather than treating disorders *per se*), such as increased positive emotions and physical activity, and stronger social connections. These are fundamental building blocks that cultivate resilience, vitality and life satisfaction and which contribute to a suite of long-term, tangible and positive physical and mental health outcomes (Alexander et al., 2021; Diener et al., 2017; Fredrickson & Joiner, 2018).

### 4.2 Nature connection exhibited a ceiling effect

Engagement in citizen science initiatives is often proposed as a means to strengthen individuals’ connection with, and concern for nature (Schuttler et al., 2018). However, our study found only slight improvements in nature connection outcomes across all five nature-based citizen science initiatives (Figure 2). This could be attributed to two factors. First, the observed ceiling effect may result from self-selection bias among participants. Given that participation was entirely voluntary, it is likely that individuals who already possess a high baseline nature connection were more inclined to participate, leaving limited scope for further improvement. Indeed, the average baseline values for NR-Self, NR-Experience and NR-Perspective were notably high (4.1, 4.0 and 4.1 respectively, on a scale of 1 – 5), surpassing the average population scores (3.29 – 3.67, 3.24 – 3.48 and 3.60 – 3.87 respectively) reported in studies from Australia and Hungary (Dean et al., 2018; Zsido et al., 2022). The potential for significant improvement might have been greater if participants had initially lower baseline scores.

Second, the relationship between people and nature is theorized to mirror personality traits, which can vary between individuals but remain relatively stable over time and across different situations (Mayer & Frantz, 2004; Nisbet et al., 2009). Consequently, substantial changes in nature connection may require prolonged and repeated exposure to nature through regular participation in citizen science initiatives. Future studies could consider two approaches – (i) adopting a randomized controlled study design to mitigate self-selection bias to more accurately determine the effects of nature exposure, notwithstanding potentially small effect sizes; and (ii) monitor longitudinal changes in nature connection through repeated assessments of individuals’ connection with nature.

Our findings on nature connection contrast recent quasi-experimental studies that report increased levels of nature connectedness following short-term engagement in citizen science initiatives. For example, Pocock et al. (2023) observed enhanced nature connection and wellbeing among 500 participants who engaged in 10-minutes citizen science and nature-noticing activities. However, it remains unclear whether these changes were due to engagement in citizen science itself, or simply from being outside the home, as the study was conducted during the 2020 COVID-19 restrictions―a period characteristic by high levels of uncertainty, stress, indoor confinement and social isolation. Similarly, White et al. (2023) reported improved nature connection and well-being among participants who interacted with birds in their gardens for at least 30 minutes, while Butler et al. (2024) found comparable for those who engaged in a 15-minute butterfly count. However, both analyses did not fully control for potential confounding effects from demographic differences, despite known variations in nature connection and well-being across factors such as age and gender (Dean et al., 2018; Oh et al., 2021).

### 4.3 Trade-offs and synergies in measured outcomes across citizen science initiatives

After controlling for potential socio-demographic confounders, we found significant improvements in positive emotions only through engagement in the QTFN initiative (Figure 3, S2). This finding was unexpected, given the well-documented associations between nature exposure and a range of health and wellbeing outcomes across different populations (Fyfe-Johnson et al., 2021; Jimenez et al., 2021). It also contrasted with the established curvilinear relationship between physical activity and health benefits, wherein notable health benefits are observed with relatively low levels of physical activity (Warburton et al., 2006). This discrepancy suggests the existence of a critical “threshold” of nature exposure, such as duration, below which health and wellbeing benefits are not realized, but above which tangible benefits become apparent. Engagement in the QTFN initiative lasted 48 hours, while that for the other initiatives ranged from 15 minutes to 4 hours.

To gain deeper insights into this threshold effect, future studies could therefore employ longitudinal study designs where the same group of participants is exposed to progressive increases in durations of engagement. However, it is essential to recognize that any identified thresholds will likely vary within and between populations (Cox et al., 2017; Oh, Fielding, Chang, et al., 2021; Shanahan et al., 2016; M. P. White et al., 2019), influenced by individuals’ baseline levels of health and nature connection, as well as the types of nature exposure and outcomes studied (Oh et al., 2021).

Second, the observed changes in health outcomes at the conclusion of the initiatives highlight the complex interplay (tradeoffs and synergies) of various components underpinning citizen science initiatives, such as the duration and intensity of engagement, and extent of social interactions. For example, a four-hour engagement through the FLOW initiative may have sufficed to improve positive emotions. However, this improvement could have been negated by the demanding nature of the task, which involved great patience and concentration to sort invertebrates into taxonomic groups.

The intensity of this engagement could have been too demanding for some participants, thereby diminishing the overall positive impact. Conversely, the QTFN initiative demonstrated significant positive changes, likely due to the synergies between the duration and social components inherent to the initiative. Participants in the QTFN initiative engaged with researchers and other volunteers over an entire weekend, in contrast to shorter durations and less-social components of other initiatives. This extended engagement provided greater opportunities for participants to apply acquired research skills in fieldwork, and to engage in problem-solving and idea exchange with a larger group. This aligns with studies highlighting the importance of social interactions within nature-based citizen science initiatives, and how participants’ satisfaction often hinge on fostering teamwork, a sense of community, and personal growth through interactions with like-minded others (Chase & Levine, 2018; Day et al., 2022).

### 4.4 Considerations for implementation

To harness the full potential of nature-based citizen science initiatives to achieve enhanced outcomes for people and biodiversity, a thoughtful consideration of participants recruitment, and the design of engagement components is essential. Citizen science initiatives like the QTFN, which balances the duration and intensity of engagement with opportunities for extended social interaction, are more likely to yield positive health outcomes in addition to achieving their primary goals. A reimagination of scientific data collection protocols, such as the duration and frequency of data collection, as measures of participants’ exposure to nature, can provide valuable insights into the health benefits derived from nature-based citizen science engagement. However, tailored approaches will be needed to assess outcomes-of-interest, and these could range from administrating pre- and post-engagement surveys, to employing holistic frameworks involving health practitioners (Skivington et al., 2021).

Achieving all scientific and non-scientific project goals to a high standard may prove challenging, but integrating social and health considerations in the goals and design of these citizen science initiatives can unlock new synergies that amplify their utility and impact. Citizen science initiatives can leverage the universal appeal of human health by elevating the health and wellbeing aspects of their recruitment strategies to attract a broader subset of society and diversify participant demographics. Collaborations with health experts can facilitate targeted evaluations of health outcomes, and the processes through which these happen, since these extend beyond the scope of citizen science coordinators. For citizen science coordinators keen to commence these evaluations, the validated scales used in our study (see Appendices A and B) provide a valuable starting point for implementation and future research endeavors. By adopting an interdisciplinary approach to evaluation and collaboration, citizen science initiatives can maximize their impact and contribute meaningfully to both scientific understanding and public health agendas.

## 5 Conclusion

Our study investigated the impact of engagement across five nature-based citizen science initiatives on eight nature connection and health and wellbeing outcomes. We observed overall improvements in all outcomes, with the greatest improvements in improved positive emotions and reduced anxiety and stress symptoms. These changes are crucial antecedents to long-term positive health outcomes. Conversely, improvements in nature connection, depression, and anxiety were modest, consistent with the view that these outcomes are less responsive to short-term interventions. The varying degrees of improvement could be attributed to several factors, including the type of participants recruited, the targeted outcomes, and the design of the citizen science initiatives, particularly the nature and social engagement components. We nonetheless advocate for reimaging citizen science initiatives as integral components of broader health-promoting interventions. By doing so, these nature-based citizen science initiatives can enhance their impact and relevance to scientific understanding, societal wellbeing and policy development.

## 6 Conflict of Interest

The authors declare that the research was conducted in the absence of any commercial or financial relationships that could be construed as a potential conflict of interest.

## 7 Funding

RO, KR, BP, SC, JG, MF and ABo acknowledge funding from the German Research Foundation (DFG-FZT 118, 202548816) that supports the German Centre for Integrative Biodiversity Research (iDiv) Halle-Jena-Leipzig. We also acknowledge funding from the Universities Australia and Deutscher Akademischer Austauschdienst (German Academic Exchange Service, DAAD) under the Australia-Germany Joint Research Co-operation Scheme, the German Ministry of Education and Research (BMBF, project grants 01BF1906 and 01UT2103A) and the Germany Agency for Nature Protection (FZ: 3520685A01).

## 8 Acknowledgments

We express our sincere gratitude to all participants in the citizen science initiatives, and to the passionate and dedicated teams leading these initiatives. We extend our thanks to past and present members from the Biodiversity and People group (UFZ/iDiv), and to our collaborators for their invaluable support, including but not limited to Franziska Lausen, Volker Grescho, Luise Ohmann, Andrea Büermann, Wayne Schmidt, Birgit Nordt and Anna Bochmann.

## 12 Data Availability Statement

The datasets analysed for this study will be made available via a freely publicly accessible repository (e.g. Zenodo) upon acceptance.

## Notes

### Competing Interest Statement

The authors have declared no competing interest.

