## Supplementary Material for "Using nature-based citizen science initiatives to enhance nature connection and mental health"

### 1 Supplementary Figures

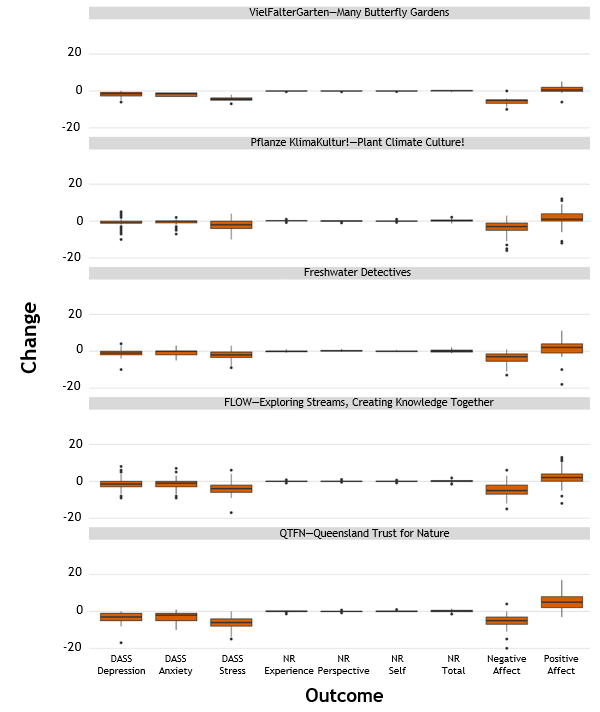

Figure S1: The change in rating for each statement was computed as the rating of statement after participation minus rating of statement before participation, for each of the five nature-based citizen science initiatives. This change could range from -4 to 4 for the dimensions of nature relatedness (NR Experience, NR Perspective, NR Self); -3 to 3 for symptoms of depression, anxiety and stress measured using the DASS-21 scale, and -4 to 4 for positive and negative affect. An improvement after exposure to nature (or engagement in nature-based citizen science initiative) was defined as a reduction in symptoms of depression, anxiety and stress (represented by negative ratings), along with fewer negative emotions (represented by negative ratings), more positive emotions (represented by positive ratings) and a stronger connection to nature (represented by positive ratings), whereas worsening signified the opposite. “NR total” was computed by summing up the change in ratings from NR Experience, NR Perspective and NR Self.

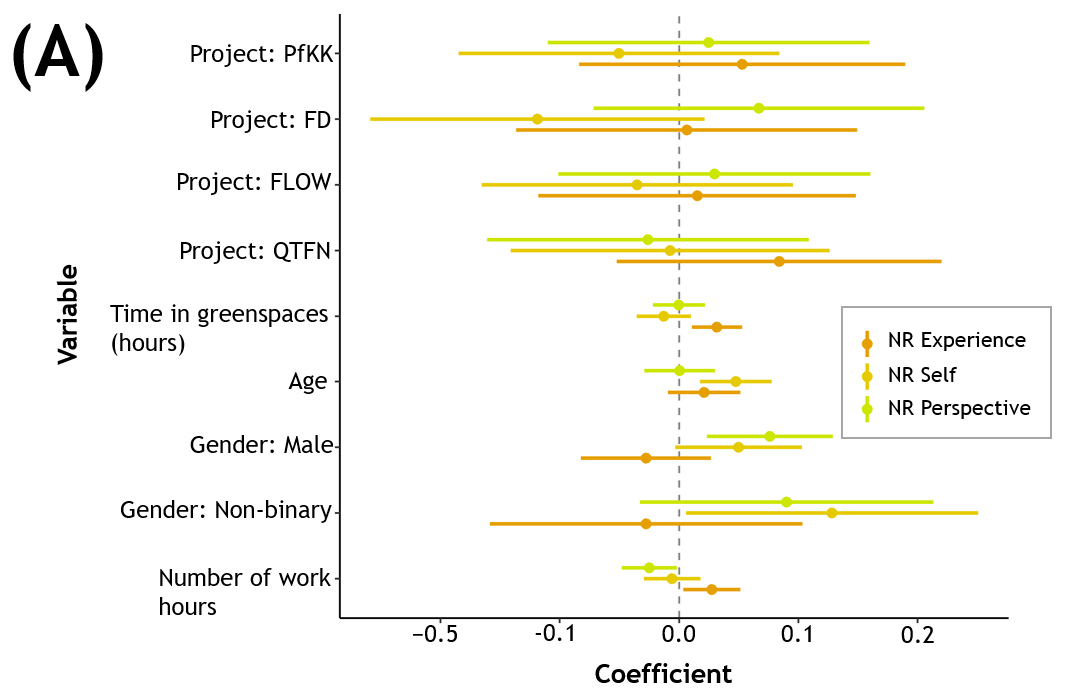

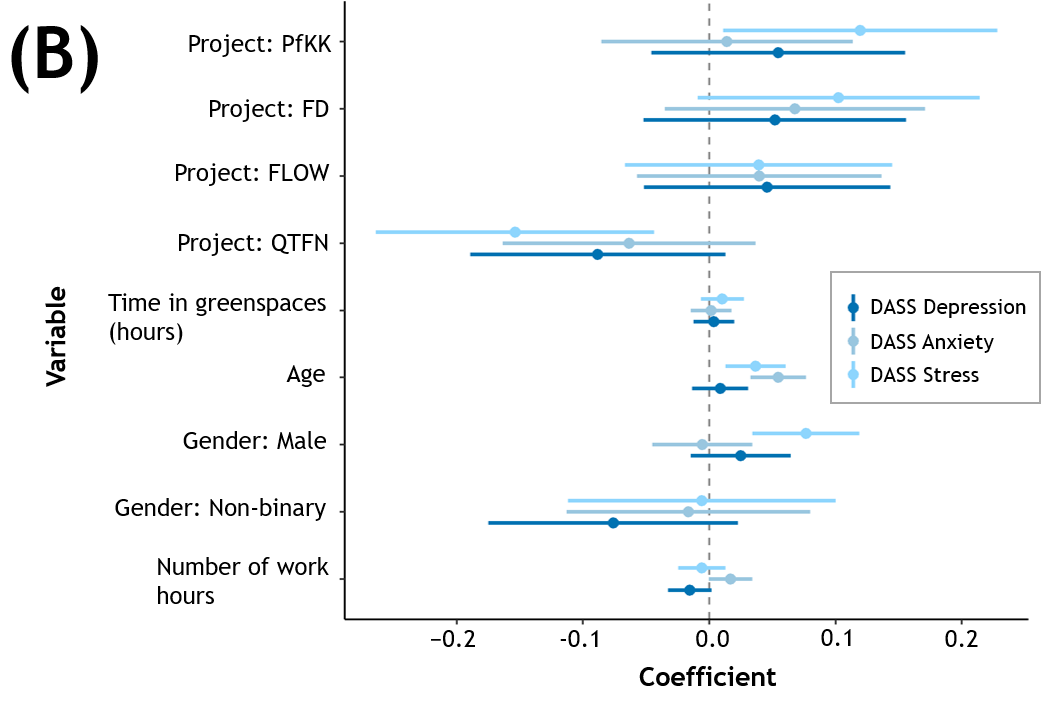

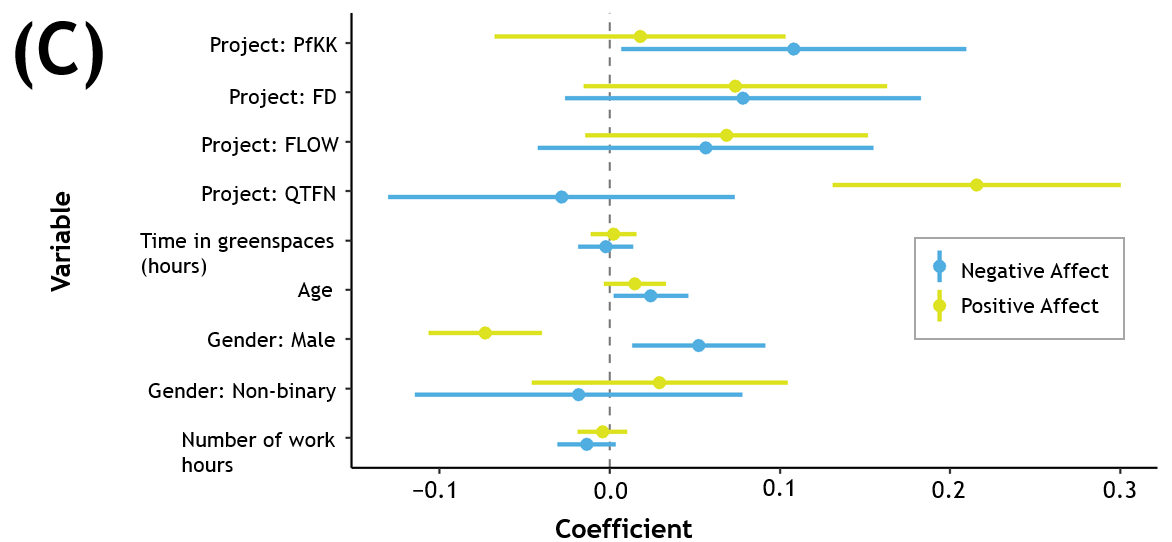

Figure S2: The model outputs from the GLMs, showing the effect size and standard errors of each predictor variable for each outcome of (a) three dimensions of nature connection (NR Experience, NR Self and NR Perspective); (b) depression, anxiety and stress as measured using the DASS-21; and (c) negative and positive affect. While we presented the effect size and standard errors for all predictor variables, only predictors Age for Anxiety, and Project: QTFN and Gender for positive affect were statistically significant coefficients (p-value ≤ 0.05). The effect size and standard errors for categorical variables are presented relative to a comparative base factor level: Project (Many Butterfly Gardens) and Gender (Female).

### 2 Appendix A – Pre-Camp Survey

Date: __________

Time: ___________

1. Please mark on the line how well you can concentrate at this moment.

Very well	Not well

2. Please mark on the line how anxious you feel at this moment.

Not at all anxious	Very anxious

3. Please mark on the line how stress you feel at this moment.

Not at all stressed	Very stressed

4. Please think about what you have been doing and experiencing during the **past week**. Then report how much you experienced each of the following feelings, using the scale below.

|  | **Very Rarely or Never** | **Rarely** | **Sometimes** | **Often** | **Very Often or Always** |
| --- | --- | --- | --- | --- | --- |
| Positive |  |  |  |  |  |
| Afraid |  |  |  |  |  |
| Good |  |  |  |  |  |
| Contented |  |  |  |  |  |
| Pleasant |  |  |  |  |  |
| Joyful |  |  |  |  |  |
| Happy |  |  |  |  |  |
| Sad |  |  |  |  |  |
| Negative |  |  |  |  |  |
| Unpleasant |  |  |  |  |  |
| Angry |  |  |  |  |  |
| Bad |  |  |  |  |  |

5. Please rate the extent to which you agree with each statement. Please select based on how you really feel, rather than how you think “most people” feel.

| **Statements** | | **Disagree Strongly** | **Disagree** | **Neutral** | **Agree** | **Agree Strongly** |
| --- | --- | --- | --- | --- | --- | --- |
| 1 | I enjoy being outdoors, even in unpleasant weather. |  |  |  |  |  |
| 2 | Some species are just meant to die out or become extinct. |  |  |  |  |  |
| 3 | Humans have the right to use natural resources any way we want. |  |  |  |  |  |
| 4 | My ideal vacation spot would be a remote wilderness area. |  |  |  |  |  |
| 5 | I always think about how my actions affect the environment. |  |  |  |  |  |
| 6 | I enjoy digging in the soil and getting my hands dirty. |  |  |  |  |  |
| 7 | My connection to nature and the environment is a part of my spirituality. |  |  |  |  |  |
| 8 | I am very aware of environmental issues. |  |  |  |  |  |
| 9 | I take notice of wildlife wherever I am. |  |  |  |  |  |
| 10 | I don’t often go out in nature. |  |  |  |  |  |
| 11 | Nothing I do will change problems in other places on the planet. |  |  |  |  |  |
| 12 | I am not separate from nature, but a part of nature. |  |  |  |  |  |
| 13 | The thought of being deep in the forest, away from civilization, is frightening. |  |  |  |  |  |
| 14 | My feelings about nature do not affect how I live my life. |  |  |  |  |  |
| 15 | Animals, birds and plants should have fewer rights than humans. |  |  |  |  |  |
| 16 | Even in the middle of the city, I notice nature around me. |  |  |  |  |  |
| 17 | My relationship to nature is an important part of who I am. |  |  |  |  |  |
| 18 | Conservation is unnecessary because nature is strong enough to recover from any human impact. |  |  |  |  |  |
| 19 | The state of non-human species is an indicator of the future for humans. |  |  |  |  |  |
| 20 | I think a lot about the suffering of animals. |  |  |  |  |  |
| 21 | I feel very connected to all living things and the earth. |  |  |  |  |  |

6. Please read each statement and select them based on how much the statement applied to you over **the past week**. There are no right or wrong answers. Do not spend too much time on any statement.

| **Statements** | | **Not applicable to me** | **Applicable some of the time** | **Applicable for a good part of time** | **Applicable for most of the time** |
| --- | --- | --- | --- | --- | --- |
| 1 | I found it hard to wind down |  |  |  |  |
| 2 | I was aware of dryness of my mouth |  |  |  |  |
| 3 | I couldn't seem to experience any positive feeling at all |  |  |  |  |
| 4 | I experienced breathing difficulty (e.g. excessively rapid breathing, breathlessness in the absence of physical exertion) |  |  |  |  |
| 5 | I found it difficult to work up the initiative to do things |  |  |  |  |
| 6 | I tended to over-react to situations |  |  |  |  |
| 7 | I experienced trembling (e.g. in the hands) |  |  |  |  |
| 8 | I felt that I was using a lot of nervous energy |  |  |  |  |
| 9 | I was worried about situations in which I might panic and make a fool of myself |  |  |  |  |
| 10 | I felt that I had nothing to look forward to |  |  |  |  |
| 11 | I found myself getting agitated |  |  |  |  |
| 12 | I found it difficult to relax |  |  |  |  |
| 13 | I felt down-hearted and blue |  |  |  |  |
| 14 | I was intolerant of anything that kept me from getting on with what I was doing |  |  |  |  |
| 15 | I felt I was close to panic |  |  |  |  |
| 16 | I was unable to become enthusiastic about anything |  |  |  |  |
| 17 | I felt I wasn't worth much as a person |  |  |  |  |
| 18 | I felt that I was rather touchy |  |  |  |  |
| 19 | I was aware of the action of my heart in the absence of physical exertion (e.g. sense of heart rate increase, heart missing a beat) |  |  |  |  |
| 20 | I felt scared without any good reason |  |  |  |  |
| 21 | I felt that life was meaningless |  |  |  |  |

7. Please create your 5-digit personal code here.

This code helps to link this survey with the follow-up survey, but all surveys will remain anonymous. cannot use it to identify persons.

Last letter of your first name, e.g. Monika = A

Last letter of your surname, e.g. Schmidt = T

Day of your birthday, e.g. 06 June 1970 = 06

First letter of your place of birth, e.g. Jena = J

CODE: AT06J

### 3 Appendix B – Post-Camp Survey

Date: __________

Time: ___________

1. Please mark on the line how well you can concentrate at this moment.

Very well	Not well

2. Please mark on the line how anxious you feel at this moment.

Not at all anxious	Very anxious

3. Please mark on the line how stress you feel at this moment.

Not at all stressed	Very stressed

4. Please think about what you have been doing and experiencing **during _____**. Then report how much you experienced each of the following feelings, using the scale below.

|  | **Very Rarely or Never** | **Rarely** | **Sometimes** | **Often** | **Very Often or Always** |
| --- | --- | --- | --- | --- | --- |
| Positive |  |  |  |  |  |
| Afraid |  |  |  |  |  |
| Good |  |  |  |  |  |
| Contented |  |  |  |  |  |
| Pleasant |  |  |  |  |  |
| Joyful |  |  |  |  |  |
| Happy |  |  |  |  |  |
| Sad |  |  |  |  |  |
| Negative |  |  |  |  |  |
| Unpleasant |  |  |  |  |  |
| Angry |  |  |  |  |  |
| Bad |  |  |  |  |  |

5. Please rate the extent to which you agree with each statement. Please select based on how you really feel, rather than how you think “most people” feel.

| **Statements** | | **Disagree Strongly** | **Disagree** | **Neutral** | **Agree** | **Agree Strongly** |
| --- | --- | --- | --- | --- | --- | --- |
| 1 | I enjoy being outdoors, even in unpleasant weather. |  |  |  |  |  |
| 2 | Some species are just meant to die out or become extinct. |  |  |  |  |  |
| 3 | Humans have the right to use natural resources any way we want. |  |  |  |  |  |
| 4 | I don’t often go out in nature. |  |  |  |  |  |
| 5 | I take notice of wildlife wherever I am. |  |  |  |  |  |
| 6 | I always think about how my actions affect the environment. |  |  |  |  |  |
| 7 | Nothing I do will change problems in other places on the planet. |  |  |  |  |  |
| 8 | I enjoy digging in the soil and getting my hands dirty. |  |  |  |  |  |
| 9 | My connection to nature and the environment is a part of my spirituality. |  |  |  |  |  |
| 10 | I am very aware of environmental issues. |  |  |  |  |  |
| 11 | My ideal vacation spot would be a remote wilderness area. |  |  |  |  |  |
| 12 | I am not separate from nature, but a part of nature. |  |  |  |  |  |
| 13 | The thought of being deep in the forest, away from civilization, is frightening. |  |  |  |  |  |
| 14 | My feelings about nature do not affect how I live my life. |  |  |  |  |  |
| 15 | Animals, birds and plants should have fewer rights than humans. |  |  |  |  |  |
| 16 | Even in the middle of the city, I notice nature around me. |  |  |  |  |  |
| 17 | My relationship to nature is an important part of who I am. |  |  |  |  |  |
| 18 | Conservation is unnecessary because nature is strong enough to recover from any human impact. |  |  |  |  |  |
| 19 | The state of non-human species is an indicator of the future for humans. |  |  |  |  |  |
| 20 | I think a lot about the suffering of animals. |  |  |  |  |  |
| 21 | I feel very connected to all living things and the earth. |  |  |  |  |  |

6. Please read each statement and select them based on how much the statement applied to you **during _____**. There are no right or wrong answers. Do not spend too much time on any statement.

| **Statements** | | **Not applicable to me** | **Applicable some of the time** | **Applicable for a good part of time** | **Applicable for most of the time** |
| --- | --- | --- | --- | --- | --- |
| 1 | I found it hard to wind down |  |  |  |  |
| 2 | I was aware of dryness of my mouth |  |  |  |  |
| 3 | I couldn't seem to experience any positive feeling at all |  |  |  |  |
| 4 | I experienced breathing difficulty (e.g. excessively rapid breathing, breathlessness in the absence of physical exertion) |  |  |  |  |
| 5 | I found it difficult to work up the initiative to do things |  |  |  |  |
| 6 | I tended to over-react to situations |  |  |  |  |
| 7 | I experienced trembling (e.g. in the hands) |  |  |  |  |
| 8 | I felt that I was using a lot of nervous energy |  |  |  |  |
| 9 | I was worried about situations in which I might panic and make a fool of myself |  |  |  |  |
| 10 | I felt that I had nothing to look forward to |  |  |  |  |
| 11 | I found myself getting agitated |  |  |  |  |
| 12 | I found it difficult to relax |  |  |  |  |
| 13 | I felt down-hearted and blue |  |  |  |  |
| 14 | I was intolerant of anything that kept me from getting on with what I was doing |  |  |  |  |
| 15 | I felt I was close to panic |  |  |  |  |
| 16 | I was unable to become enthusiastic about anything |  |  |  |  |
| 17 | I felt I wasn't worth much as a person |  |  |  |  |
| 18 | I felt that I was rather touchy |  |  |  |  |
| 19 | I was aware of the action of my heart in the absence of physical exertion (e.g. sense of heart rate increase, heart missing a beat) |  |  |  |  |
| 20 | I felt scared without any good reason |  |  |  |  |
| 21 | I felt that life was meaningless |  |  |  |  |

7. **About how often do you usually visit or pass through outdoor greenspaces for any reason?** *For example, this includes beaches, picnic areas, children's playgrounds, national parks, golf courses, tennis courts, bike-ways and bowling greens.*

- 6-7 days a week
- 3-5 days a week
- 2-3 days a week
- Once a week
- 2-3 times a month
- Once a month
- Once every three months
- Once a year
- Never

8. **Over the last week, what outdoor greenspaces did you visit or pass through? Can you estimate the total time you spent there?** *This includes, for example, beaches, children's playgrounds, parks, bushland, bike-ways, picnic areas, beaches, golf courses, tennis courts and bowling greens. Note: Please write in number of hours. if you did not spend any time, please indicate 0. If you spend 30 min, please indicate 0.5 hours. If you spend 1 hour, please indicate 1.*

*___________ Hours*

9. Please rate the extent to which you agree with each statement.

| **Statements** | | **Disagree Strongly** | **Disagree** | **Neutral** | **Agree** | **Agree Strongly** |
| --- | --- | --- | --- | --- | --- | --- |
| 1 | My friends spend a lot of time in nature |  |  |  |  |  |
| 2 | My family approve of me spending time in nature |  |  |  |  |  |
| 3 | I am very close to my family |  |  |  |  |  |
| 4 | My friends approve of me spending time in nature |  |  |  |  |  |
| 5 | My family spends a lot of time in nature |  |  |  |  |  |
| 6 | I am very close to my friends |  |  |  |  |  |

10. Please rate your own health.

| Very poor | Poor | Average | Good | Very good |
| --- | --- | --- | --- | --- |
| 1 | 2 | 3 | 4 | 5 |

8. **What is the postal code of where you currently live?** (We would like to use this information to calculate the percentage of green space around your residence. All information will be anonymized, and presented as a summary only).

11. How old are you?

_______ years old

12. What is your gender?

- Female
- Male
- Non-binary

13. Are you now in paid employment?

- Yes, working
- No, not working but looking for job
- Home duties
- Don’t work
- Retired
- Student

14. If you are working, approximately how many hours do you spend at work in a normal week?

- No time
- 5 hours or less
- 6-10 hours
- 11-20 hours
- 31-40 hours
- 41-50 hours
- 51-60 hours
- 61+ hours

15. What is the level of the highest qualification or schooling year you have completed? (please tick one)

- Post-graduate degree
- Associate / Advanced diploma
- Bachelor degree
- Trade Certificate
- Year 9 - 12 or equivalent
- Year 8 or below
- Other (please specify)________________________________________

16. Please create your 5-digit personal code here.

This code helps to link this survey with the follow-up survey, but all surveys will remain anonymous. cannot use it to identify persons.

Last letter of your first name, e.g. Monika = A

Last letter of your surname, e.g. Schmidt = T

Day of your birthday, e.g. 06 June 1970 = 06

First letter of your place of birth, e.g. Jena = J

CODE: AT06J
